# Neuro-musculoskeletal Upper Limb in-silico as virtual patient

**DOI:** 10.1101/2021.05.16.444298

**Authors:** Mallampalli Kapardi, Madhav Vinodh Pithapuram, Yashaswini Mandayam Rangayyan, Raghu Sesha Iyengar, Avinash Kumar Singh, Sirisha Sripada, Mohan Raghavan

**Author notes:** The authors would like to gratefully acknowledge the support by way of grant-in-aid provided by the Ministry of Electronics and Information Technology(MEITy-4(11)/2018-ITEA), Government of India.

## Abstract

Virtual patients and physiologies allow experimentation, design, and early-stage clinical trials in-silico. Virtual patient technology for human movement systems that encompasses musculoskeleton and its neural control are few and far in between. In this work, we present one such neuro-musculoskeletal upper limb in-silico model. This upper limb is both modular in architecture and generates movement as an emergent phenomenon out of a multiscale co-simulation of spinal cord neural control and musculoskeletal dynamics. It is developed on the NEUROiD movement simulation platform that enables a co-simulation of popular neural simulator NEURON and the musculoskeletal simulator OpenSim. In this work, we describe the design and development of the upper limb in a modular fashion, while reusing existing models and modules. We further characterize and demonstrate the use of this model in generating a range of commonly observed movements by means of a spatio temporal stimulation pattern delivered to the cervical spinal cord. We believe this work enables a first and small step towards an in-silico paradigms for understanding upper limb movement, disease pathology, medication, and rehabilitation.

## I. Introduction

Virtual patient is defined as an interactive computer simulation program of real-time clinical scenarios for assessment, education, and medical training [1]. The development of virtual patients has revolutionized the healthcare system globally. They provide us an opportunity to understand complex functions, integrate relevant physiological and anatomical data, accelerate the development of therapies and medical devices [2]. An essential feature of virtual patients is the simulation of a particular clinical outcome of interest. This also implies that the relevant physiological manifestations, their external symptoms, and the internal variables be represented in the model too. Virtual patients must also incorporate the patient cohort variation in human anatomy/physiology and other statistical variations [3], [4]. Currently, most of the virtual patients are focused on cardiac physiology and orthopedics [5]. In today’s scenario, orthopedic simulations focus largely on the biomechanics of the musculoskeletal system with little or no neural components built-in [6]. Motor systems neuroscience especially deals with the final common pathway - the spinal cord which had been known and understood for a long time [7] and hence presents the lowest hanging fruit in the pursuit of virtual movement.

The motor commands come from the higher centers of the brain, the frontal and posterior parietal cortex (PPC). Areas like the premotor and supplementary motor cortex help in planning an action sequence. But, they require basal ganglia (BG) to give input. BG has two types of connections, firstly a direct connection that selects a particular action to initiate and the latter indirect connection which disregards the unnecessary motor commands [8]. According to the literature the layer V of M1 contains numerous pyramidal neurons that connect to the spinal cord through the corticospinal tract. It is the axons of these pyramidal neurons that are connected to the motor neurons in the spinal cord monosynaptically to activate muscle fibers [8]. The primary motor cortex (M1) is responsible for the execution of motor programs at lower levels [9]. Motor programs may be thought as a pattern of time-varying input to various spinal neurons that results in a specific movement such as center-out movement at various angles. The BG influence the choice of motor programs in a given context.

The cervical segments (C3 - C8) and the initial thoracic segment (T1) are known to be active for most of the upper limb muscle movements, which consist of shoulder to digit movements [10]. Motor neurons in the ventral part of spinal cord segments and neurons in dorsal root ganglion (DRG) are the prime connectors between the neural and musculoskeletal systems [11], [12]. The motor neurons directly activate the muscle fibers whereas the DRG act as sensors of the musculoskeletal system in form of proprioception to control and deliver a proper kinesthetic response to the muscles [13]. Proprioception plays an important role in sensorimotor integration [14]. The interneurons act as local regulators for handling muscle synergies in the musculoskeletal system [15]. Various models of the motor neurons, interneurons, and Renshaw cells were created to understand and investigate the ion channel properties, complex firing patterns, effects of soma and dendritic distributions [11], [12], [16], [17]. All these properties vary vastly with motor units and spinal cord segments.

Biomechanical models of the musculoskeletal systems have helped in understanding the neuromuscular control, muscle moment arms, and to design prosthetics [18], [19]. There are many models designed for the upper limb as well, which consist of all upper limb muscles. The models also include ribs, scapula, humerus, radius, ulna, bones of the wrist, and hand [20], [21]. These models have couplers between joints which are used extensively for understanding the influence of multiple degrees of freedom over the other joints. In addition, the force generation characteristics of the muscles in the model are determined using the Hill-type muscle model, which requires at least four parameters namely pennation angle, peak force, optimal fiber length, and tendon slack length.

While there is a lot known about movement infrastructure and its mechanisms, most current models of movement are restricted to one or the other domains of neural [22]–[24] or musculoskeletal domains [25]. The few models that build neuro-musculoskeletal models are not modular, generalizable [26], [27], and most importantly do not incorporate anatomical features of the spinal cord. The NEUROiD movement platform [28] has made it possible to create movement infrastructure by combining available musculoskeletal and spinal circuit models in a modular fashion within an anatomical precinct. In our paper, we propose to build a virtual upper limb by curating well known spinal circuits located within spinal cord anatomical regions and combine the same with published musculoskeletal models of the upper limb. We demonstrate broad similarities with intraspinal stimulation studies [29]. We also use the model to understand the complex interplay between the spinal neural circuits and the constraints of the musculoskeletal system and how they influence each other in a close loop environment to produce movement.

## II. Methods

The upper limb model had two entities a musculoskeletal model simulated using OpenSim and a spinal cord model simulated on NEURON. Development of neuro-musculoskeletal model took place on NEUROiD platform [28]. NEUROiD’s model builder encapsulated the spinal cord model and the OpenSim model. Fig. 1(a) show casts the NEUROiD interface.

**Fig. 1.**
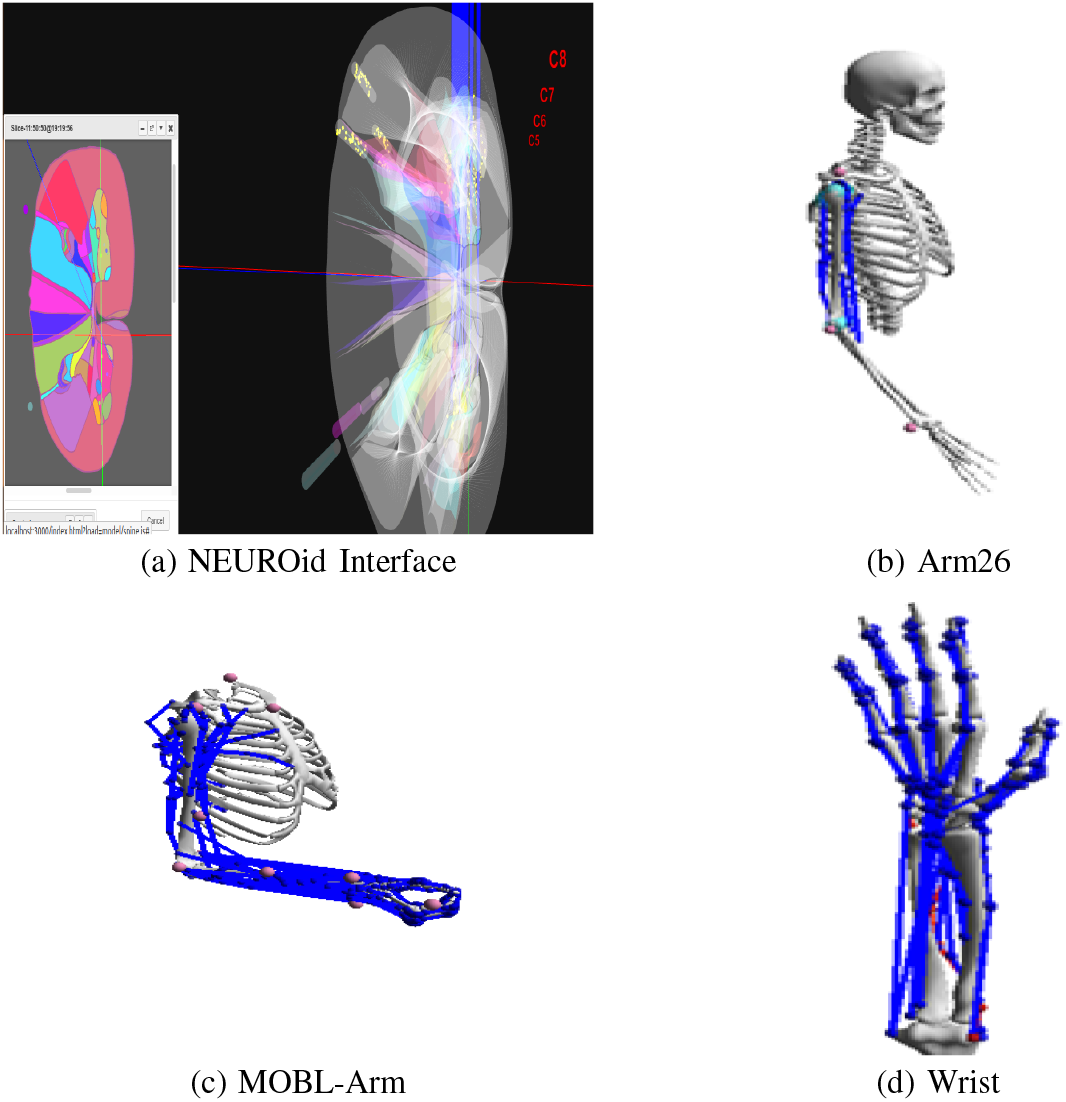
Embodies NEUROiD and different OpenSim Models facilitated, namely Arm26, MOBL-Arm and wrist model. The Fig 1(a) depicts the anatomy of spinal cord from C4-T1 segments in rostro-caudal direction looking in from rostral side. Blue electrodes are for recording output responses and red electrodes for input stimulation. Yellow circles represent the cell groups. Arm26 is a simple three muscle model with a single active joint, MOBL-Arm is sophisticated model containing fifty muscles and 7 degrees of freedom. Wrist model consists of 25 muscles controlling the movement of all the digits in the model.

### A. Spinal Cord model and circuits

The spinal circuits responsible for the movement of upper extreme muscles are found in cervical and thoracic segments. Spinal cord segments from C4-T1 were considered in our model for the simulation. The motor circuit primarily consists of interconnections between alpha motor neurons, interneurons, Renshaw cells, and proprioceptive afferents (Ia, Ib, II). The cell groups and connections are specified hierarchically and in a rule based manner within NEUROiD’s model building framework. Table I demonstrates cell specification as a combination of functional cell type (e.g., alpha motor neurons, interneurons, afferents) and the muscle to which it is related. The locations of cell groups in particular lamina in each spinal cord segment were based on [30], [31]. The motor neuron cell types are based on [32] while other neuron types are modeled as simple Hodgkin Huxley neuron models.

**TABLE I.**
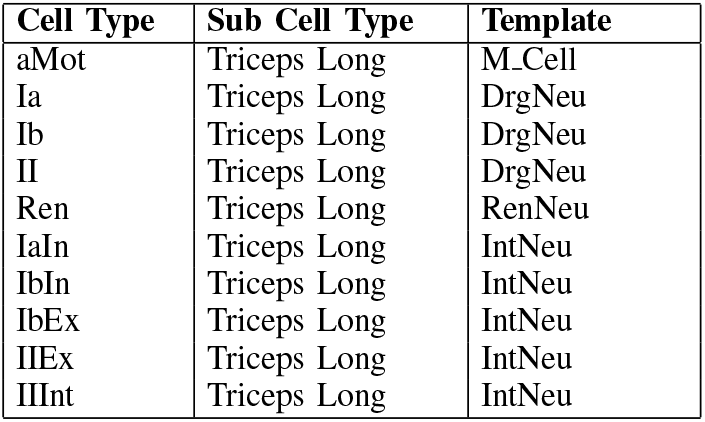
Cell Template Map

The connections between the neurons are also specified as a set of rules in a spread sheet. The connections are based on existing literature, where Ia DRG neurons connect monosynaptically to the alpha motor neuron of the homonymous muscles and other agonist muscles through their respective interneurons [33]. The Ib afferents excite the antagonistic motor neurons via interneurons, while inhibiting homonymous muscle motor neurons through interneuron relays. The group II afferents excite the motor neuron of the homonymous and synergist muscle and inhibits the corresponding antagonist muscle [33]. The Renshaw cells perform gain control in the circuit by inhibiting the motor neuron of homonymous muscle and also inhibit the Ia interneuron of antagonist muscle [34].

The connections rules are tabulated in Table II. A typical rule mentions the pre-synaptic, post synaptic neuron types (e.g., motor neurons, interneurons, afferents) and the muscles to which the rule must be applied (self, homonymous, antagonist, ipsilateral or contralateral). In addition, the rules also specify the kind of synapse (excitatory or inhibitory) and the strength of the connection. The convergence within the cell groups is set to 0.5 and conduction delay for Ia afferent is 15 ms, for Ib and II afferents is 30 ms. The activation of a muscle is in the range [0,1] and considered to be directly proportional to the neural activity of its motor neurons, where maximal motor neuron activation u(t) is 1.

**TABLE II.** Connection Rules

| Connection type | Source | Target | Target Muscle group | Strength and Type of Synapses |
| --- | --- | --- | --- | --- |
| Ia-excitation | Ia | aMot | self,agonist | 0.6, EXC |
| Inter-moto | Ia | IaIn | self | 0.53, EXC |
| Inter-moto | Ib | IbEx | self | 0.55, EXC |
| Inter-moto | Ib | IbIn | self, agonist | 0.59, EXC |
| II-excitation | IIEx | aMot | self | 0.6, EXC |
| II-inhibition | IIInt | aMot | antagonist | 0.5, IHB |
| Inter-moto | II | IIEx | self | 0.5, EXC |
| Inter-moto | II | IIInt | self | 0.5, EXC |
| Ia-inhibition | IaIn | aMot | antagonist | 0.52, IHB |
| Ib-inhibition | IbIn | aMot | self | 0.58, IHB |
| Ib-excitation | IbEx | aMot | antagonist | 0.65, IHB |
| Renshaw-activation | aMot | Ren | self, agonist | 0.6, EXC |
| Renshaw-moto | Ren | aMot | self | 0.54, EXC |
| Reciprocal-inhibition | Ren | IaIn | antagonist | -0.7, IHB |

EXC represents excitatory synapse and IHB represents inhibitory synapse. The decay time constant for excitatory and inhibitory synapses are 0.75 ms and 1.5 ms respectively. The reversal potential values for excitatory and inhibitory synapses are 0 mV and -80 mV.

**TABLE III.**
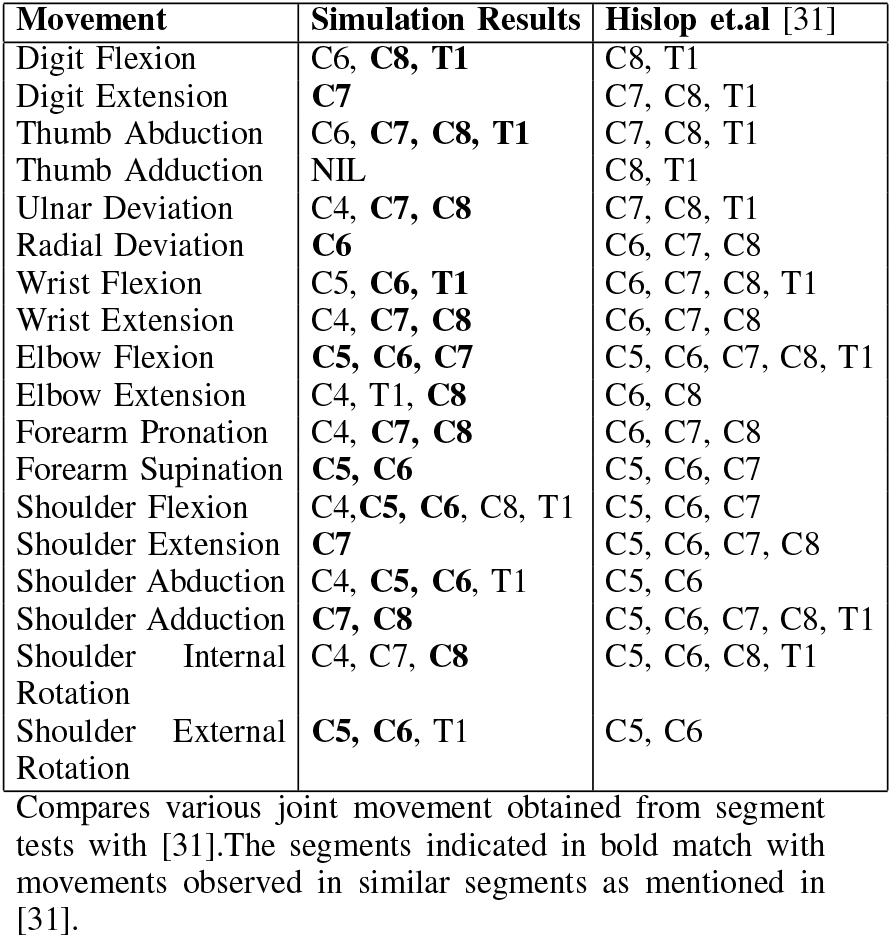
Validation of the model

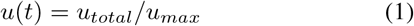

Where u(t), is motor neuron excitation, *u*_*total*_ is total firing frequency of the motor neuron group, *u*_*max*_ is the maximum firing frequency of the motor neuron.

The firing rates of proprioceptive feedback was calculated based on [35].

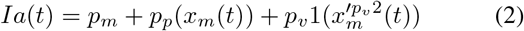

where *x*_*m*_(*t*) is muscle stretch in mm,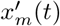 is muscle stretch velocity in mm/s. pp is length change constant and velocity constants are *p*_*v*1_ and *p*_*v*2_. *p*_*p*_ = 13.5, *p*_*v*1_= 4.3, *p*_*v*2_ = 0.6.

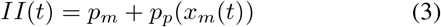

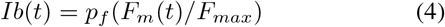

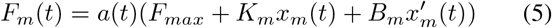

Where, *p*_*m*_ = 5 sp/s, *p*_*f*_ = 200 sp/s, muscle stiffness *K*_*m*_ = 56.3 kN/m, damping *B*_*m*_ = 2.81 kNs/m, *F*_*max*_ is the maximum isometric force of the muscle, *F*_*m*_(*t*) is muscle force in N, a(t) is the muscle activation.

### B. Musculoskeletal Models

Three OpenSim models were considered for modeling and understanding the dynamics of upper limb muscles. The Arm26 model Fig. 1(a) consists of 6 muscles namely Biceps (2 muscle heads), Triceps (3 muscle heads), and Brachialis [20]. The Arm26 model was constrained by shoulder joint.The MOBL-Arm model Fig. 1(b) consists of 50 muscles that include shoulder, elbow, and wrist joints, with 15 degrees of freedom. However it does not support any digit movements and does not have neck muscles to support shoulder movement efficiently [20]. The third model Fig. 1(c) is a wrist-only model and consisted of 15 muscles. The kinematics of the wrist model was constrained by wrist flexion-extension [21].

The single joint model (Arm26) was used to test and characterize the efferent and afferent interfaces between the spinal cord and musculoskeleton. Multi-joint movements were tested on MOBL-Arms model. Wrist model was used to exercise the ability of the spinal cord to manipulate the digits. The initial states of all the OpenSim models were chosen such that there are no residual forces to start with and the limb is in equilibrium with no application of stimulation to the spinal cord. Sanity tests were performed to verify the integrity of all muscle-motor neuron connections and the ability of the model to exercise all the relevant degrees of freedom available. Further tests were performed to characterize the effects of some important model parameters where the flexors and extensors were stimulated separately with and without the afferent in closed loop in an Arm26 model. This was done on multiple model variants with different proprioceptive afferents Ia, Ib and II. Similar experiments were also conducted to evaluate the effect of afferent synaptic weights, high (HW: 0.8-1.0), average (NW: 0.5-0.75) and low weights (LW: 0.1-0.25).

### C. Individual Segment Test

This mimics the spinal cord electrical stimulation experiments that map the range of upper limb movements onto the spinal cord segments [29]. However, as detailed information of the specific spinal neuron groups stimulated in each of the micro-stimulation experiments were not available, we perform a single stimulation experiment per spinal segment that stimulates all the motor neurons present in the segment. The motor neuron groups were stimulated by intracellular current injections in the distal end of dendrite with a square pulse of width 200 ms and amplitude of 85 nA current, that induced an approximately 60 Hz spiking in the soma of alpha motor neuron. The MOBL-Arm model was stimulated at a similar spiking frequency (60 Hz), while the wrist was stimulated with a lower frequency (20 Hz). This is in line with the relative stimulation currents used in [29]. Total simulation time for experiments involving Arm26 and wrist model was 200 ms and for MOBL-Arm it was 300 ms. For each stimulation experiment we observed the range of motion elicited in each joint and each degree of freedom therein. The segment tests were conducted for all segments from the fourth cervical segment (C4) to the first thoracic segment (T1) [10]. These tests were performed for all the three musculoskeletal models and in two configurations: open loop (with afferent proprioceptive fibres severed) and closed loop (intact proprioceptive afferents)

### D. All Segment Test

While the previous tests observed the movement types generated by a constant stimulus pattern, it is essential to understand how the model responds to a sequence of different stimuli, possibly generating antagonistic movement types. For the model to be usable for movement simulation, the model must be able to generate a smooth movement in response to an input stimulus that changes in continuous or discrete steps. In order to test the ability of the model to generate smooth movements all the spinal cord segments (their motor neurons) were stimulated one after another in ascending and descending orders (T1 - C4 and C4 - T1). Stimulation of each segment was for a duration of 50 ms at 60Hz. The musculoskeletal model used for this test was MOBL-Arm. The total simulation time was 600 ms.

### E. Varying Initial Postures Test

Different postures represent distinct states of the musculoskeletal model with their own mechanical advantages. In order to evaluate the suitability of our model to evaluate these properties, we performed experiments with identical electrical stimulation patterns but with different initial postures of the musculoskeletal model. We chose a stimulation protocol similar that used for characterizing afferents. The upper limb was tested with elbow in three different positions (90, 60 and 30 degrees). Resultant movements obtained were analyzed and correlated with known results.

## III. Results

Spinal cord anatomy and physiology were integrated with several models of upper limb musculoskeleton to create *in-silico* models. In each case, the constituent musculoskeletal models were exercised by direct stimulation from native OpenSim graphical user interfaces for all the joint movements in varying degrees of freedom. Similarly, the integrity of neuromuscular interfaces were tested by targeted stimulation of various spinal motor neuron cell groups and their ability to contract the respective muscles were verified.

### A. Model Characterization

#### 1) Effect of Afferents

Experiments were performed on model versions with all proprioceptive afferents turned off (Open Loop), only one of Ia, Ib or type II afferents turned on. The corresponding results are shown in Fig. 2. The Open loop responses (OL) to stimulation of flexor or extensor groups represents the inherent ability of muscle groups to effect flexion or extension, starting from a given initial position. In this case it may be seen that for a given stimulation strength and a starting position of 60 degrees flexion of elbow joint, the flexors achieve a larger range of motion than the extensors. It may also be seen that Ia and Ib afferents resist the movement and reduced the range of motion compared to OL, which is consistent with standard results in physiology [36]. The resistance of the type II afferents are largely dependant on the amount of stretch from optimal muscle lengths [37]. Since in this experiment we start with a slightly flexed position, the extensors are already stretched. As a result, the type II afferents tend to favour extension.

**Fig. 2.**
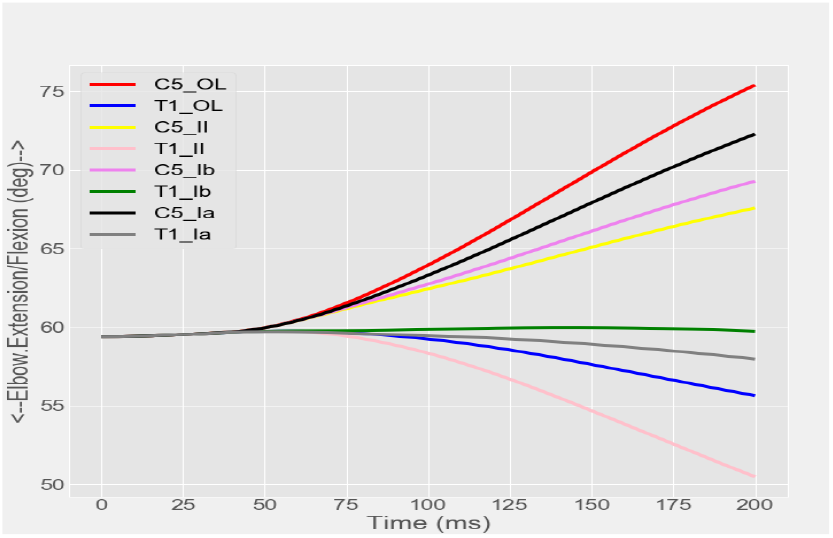
Effects of each afferent were tested in the model by stimulating biceps, brachialis related alpha motor neurons in C5 segment and triceps related motor neurons in T1 segment at 60 Hz for 100 ms. Both Ia and Ib afferents damped the flexion or extension of the elbow joint. Type II afferents exaggerated extension and strongly resisted flexion.

#### 2) Effect of differential afferent delay with multiple afferents

When all the proprioceptive afferents are at play simultaneously (Closed Loop - CL), their relative peripheral latencies determine their contributions to the model responses. Experiments suggest that the latencies of Ia are around 15 ms while the Ib and type II latencies are slightly larger at about 30 ms. Under these circumstances it may be seen Fig. 3 that CL almost follows the Ia response. However as the gap in latencies between the afferents narrow down, the CL responses strike a balance between the responses of Ia, Ib and II afferents.

**Fig. 3.**
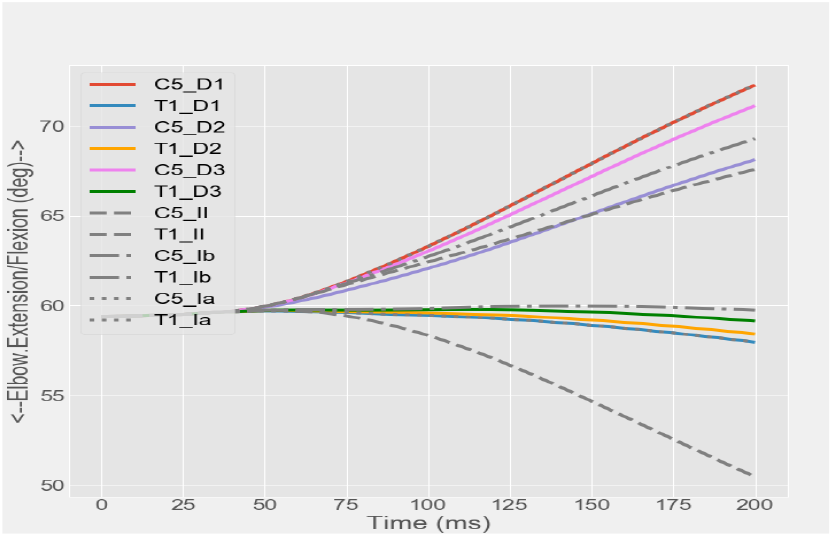
Effects of conduction delay was tested in the model by stimulating biceps, brachialis related alpha motor neurons in C5 segment and triceps related motor neurons in T1 segment at 60 Hz for 100 ms. Three conditions were considered where the conduction delay of Ia and other afferents were set respectively to 15 and 30 ms (D1); 19 and 26 ms (D2); 22.5 and 22.5 ms (D3). Shorter delays significantly increased the influence of the afferent type in the composite circuit consisting of multiple afferents. The joint trajectory in presence of single afferents (Ia, Ib, II) are indicated in gray lines for reference. It may be seen that for very short Ia delays (D1) the joint trajectory mirrors Ia. However as delays of afferents are comparable (D2) or equal (D3) the joint trajectory observed is increasingly influenced by multiple afferents.

#### 3) Effect of Initial Postures

Effect of initial posture of the model plays a crucial role in the resulting behavior of the model. Three different joint positions were considered to test the effect of postures on elbow joint - elbow flexion of 30, 60 and 90 degrees. It may be seen in Fig. 4 that although the stimulus to the spinal cord is identical, the initial musculoskeletal position alters the flow of proprioceptive afferents and thereby alters the behavioural (movement) response. It may be seen that maximum range of extension was seen when starting with a more flexed position, and flexion was maximized when starting with less flexed positions. This is in conformance with known behaviour of proprioceptive afferents that try to move muscles away from excessively stretched or compressed states towards their optimal lengths [36].

**Fig. 4.**
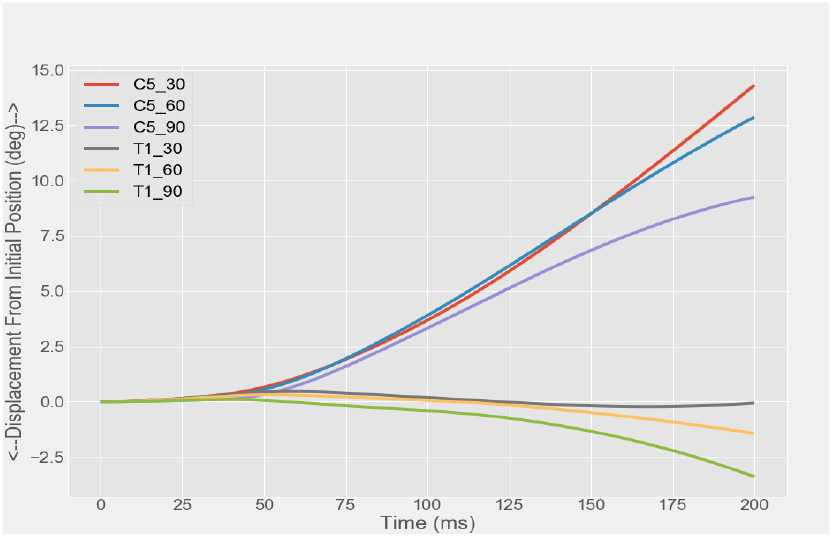
Initial state of a joint effects the behavioral outcome of the biomechanical model. Biceps, Brachialis related alpha motor neurons in C5 segment and triceps related motor neurons in T1 segment were stimulated at 60 Hz for 100 ms. Three different states were considered where the elbow joint was placed at 90,60 and 30 degrees respectively. It may be seen that resistance to flexion and exaggerated extensions are seen when starting from increasingly flexed positions.

#### 4) Connection Strength

Needless to say that connection strengths play a role in deciding the dominance of one type of afferent over the other when multiple afferents are active. However, experimental measurements of their relative synaptic strengths are not known. The Fig. 5 shows that increasing afferent synaptic strength increased the amount of resistance to movement compared to OL. Extremely high afferent synaptic strength can lead to unnatural results such as flexion when extensors are excited and vice versa.

**Fig. 5.**
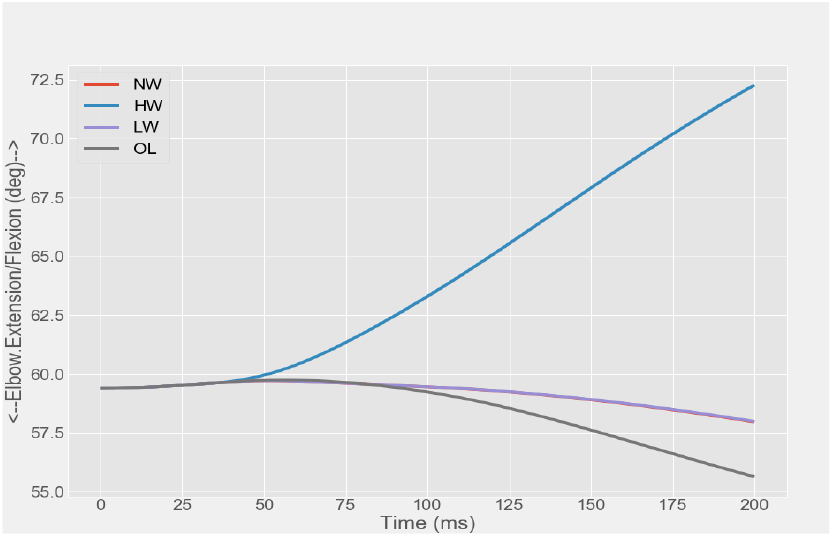
Effects of connection strengths in motor circuit were analysed. Triceps related alpha motor neurons in T1 segment were stimulated at 60 Hz for 100 ms. Increased synaptic strengths of afferent result in increased resistance to motion. Extremely high weights can even invert the type of movement.

### B. Validation of the model from Segment Tests

Simulations of segment wise stimulation resulted in the emergence of a variety of joint movements across different segments in the full arm model Fig. 6 and wrist model Fig. 7. The videos of limb movement corresponding to these simulations are provided as part of supplementary information. The broad distribution of these different movement types were compared with that of [31]. Since we use a single stimulation covering all motor neuron groups in a segment, in our experiments every cervical segment can only be associated with a single type of movement (per degree of freedom), and most likely with the movement type mediated by the dominant and numerous motor neuron group. However within these limitations, broad agreements between our results and that of [31] were found in Table II. Comparisons with micro stimulation experiments of [29] also showed that there was broad agreement. However a more rigorous analysis could not be performed since the specific cell groups recruited in each micro stimulation of [29] was not known and they were performed on mice.

**Fig. 6.**
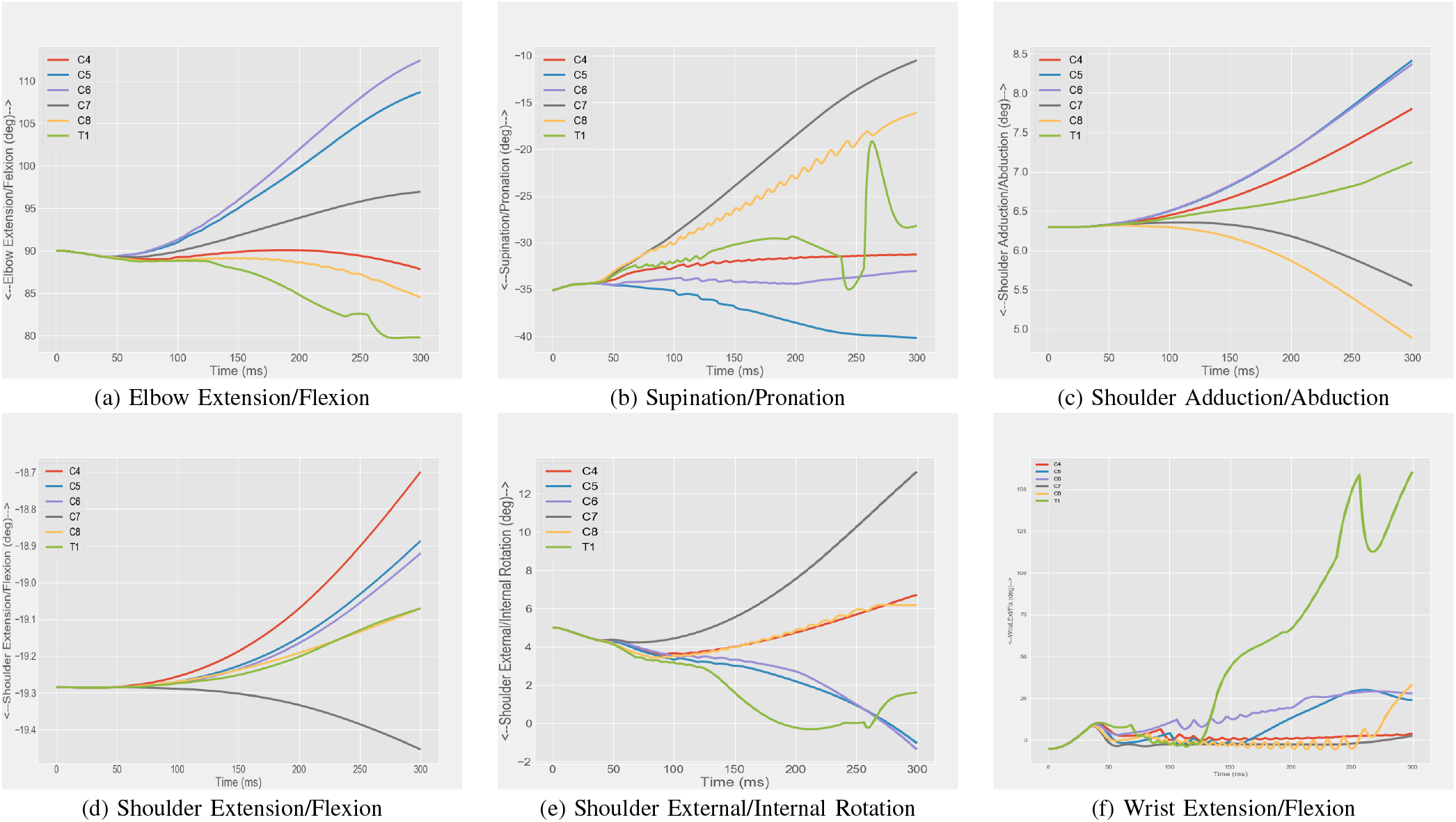
Segment test from C4 to T1 was performed for MOBL-Arm model. Every segment was stimulated at 60 Hz for 200 ms. The elbow joint was flexed to 90 degrees and wrist was deviated 18 degrees radially before the start of simulation. Variations in major joint angles and their degrees of freedom are depicted in the graph.

**Fig. 7.**
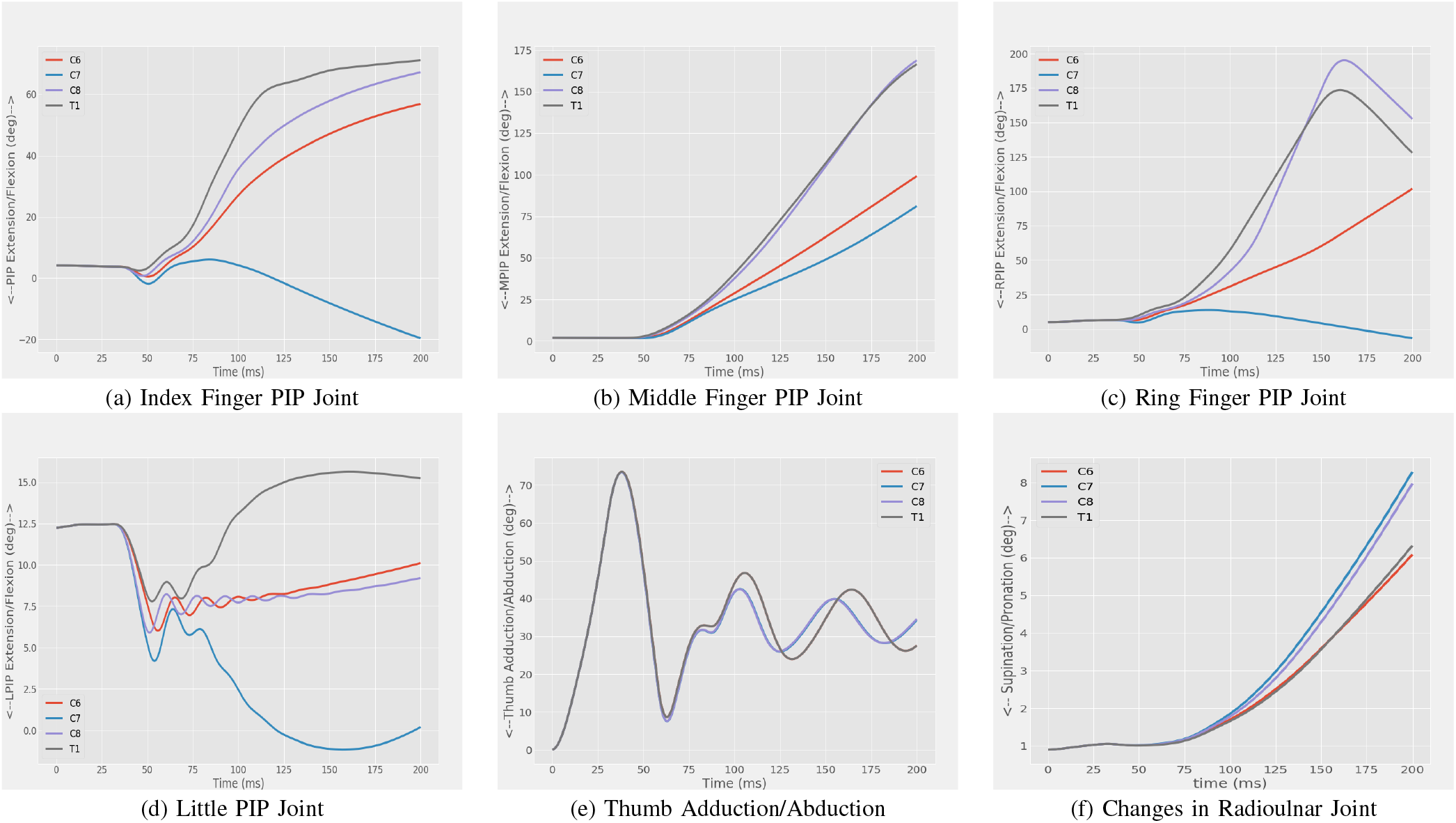
Segment test from C6 to T1 was performed for Wrist model. Every segment was stimulated at 20 Hz for 100 ms. The proximal interphalangeal joint of all digits in Wrist Model were compared. The simulations resulted in flexion of all digits, during the stimulation of C7 segment the little, ring and index digits had an extension of proximal interphalangeal joint (PIP) joint.

### C. All segment Test

The all segment tests showed at Fig. 8 that the model was capable of executing a smooth movement in response to a sequence of spatio-temporal stimulation of the cervical spinal cord. The set of movements executed in sequence quite often consisted of antagonistic movements like flexion/extension and supination/pronation. The corresponding movements were smooth to visual inspection (Supplementary material). The transitions between antagonistic movement types was smooth in the shoulder and elbow joints as seen from the graphs of joint angles as well. The wrist joint exhibited some jerkiness compared the shoulder and elbow joints. This is again consistent with published results [38] and the fact that the stimulation in our experiment was a segment wise stimulation without any fine targeting.

**Fig. 8.**
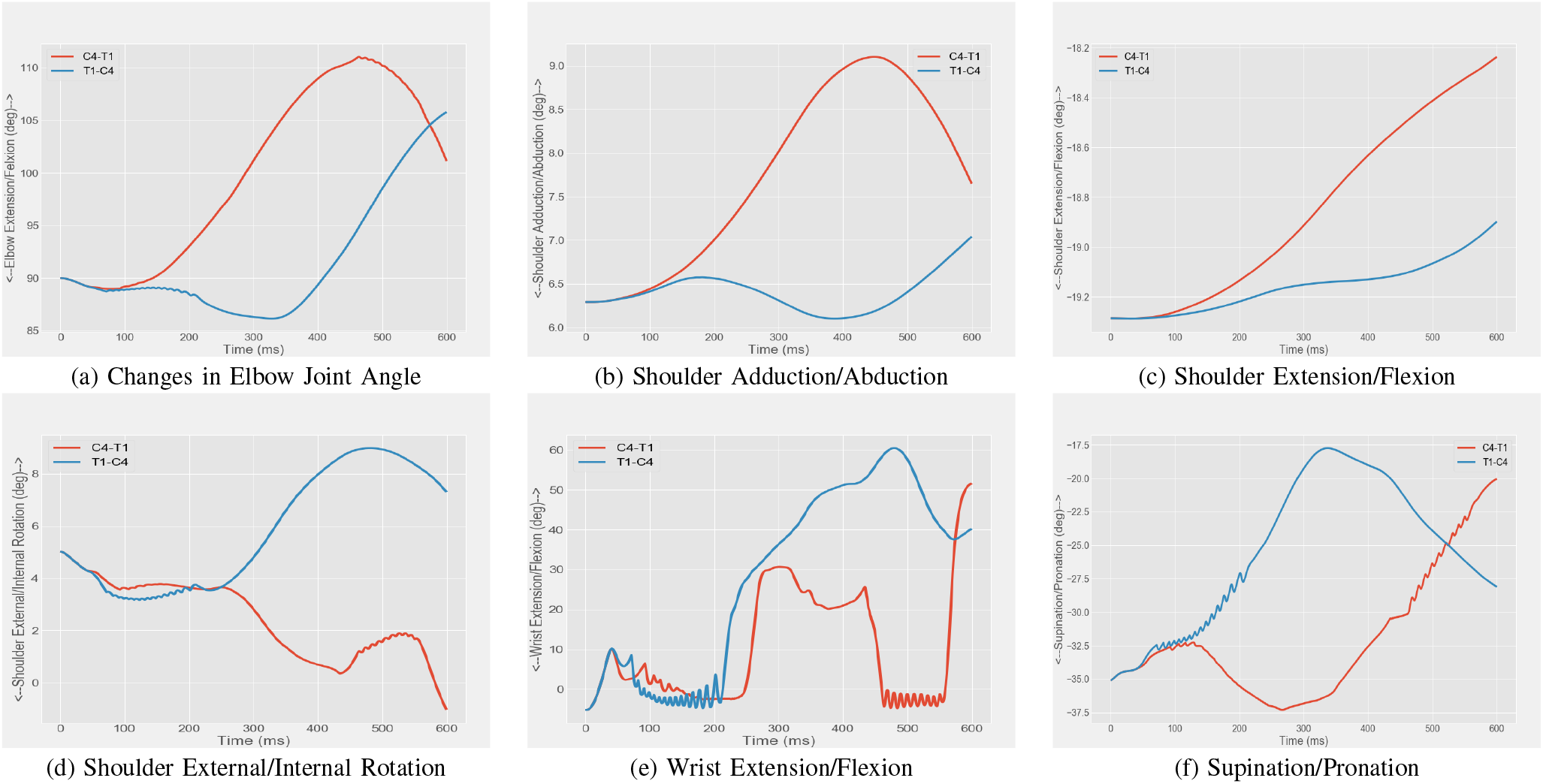
The all segment test (AST) each segment was stimulated at 60 Hz for 50 ms, the delay between each stimulation of segment was 50 ms. The elbow joint was flexed to 90 degrees and wrist was deviated 18 degrees radially before the start of simulation. The joint angles for each degree of freedom are compared for C4-T1 and T1-C4 tests.

## IV. Discussion

We have demonstrated a first step towards a full in-silico neuro-musculoskeletal upper limb by building a spino-musculoskeletal upper limb in-silico, using the NEUROiD neuro-musculoskeletal co-simulation platform. Our upper limb model in-silico integrates a variety of anatomical, physiological data gleaned by experimentation with regard to the nervous and musculoskeletal systems. The neural model itself is multiscale with representations of ion-channel, neuron, synapse, cell groups and pathways stitched together hierarchically and tied to relevant anatomical locations of the cervical spinal cord. In such an in-silico upper limb, we demonstrate the emergence of movement as an emergent phenomenon, as a result of focal electrical stimulation of cervical spinal cord. We also demonstrated the broad similarity of responses in our in-silico model to spinal cord micro-stimulation experiments in terms of different movement types elicited by stimulation at various spinal cord segments. We also characterize the effect of specific afferent types, connection strengths and musculoskeletal postures on the elicited movements. Our upper limb is capable of continuous movement in response to a spatio-temporal spinal cord stimulation pattern as would be expected physiologically in vivo. Our limb is modular and easily supports replacement of components at any scale from cellular to system. We also demonstrate the inter-operability of our cervical spinal cord model designed in NEUROiD with three different musculoskeletal models in OpenSim. While this model implements neural elements only up to the level of cervical spinal cord, interfacing the same with upstream models of the cortex, BG or cerebellum using the popular NEURON neural simulator are straightforward. While all our model parameters are based on published literature, it must be emphasized that debates on the suitability or otherwise of model parameters are possible, as they must be. Further synaptic weight values are not available in published literature and parameters are chosen to be values that result in measured movement behaviour [29]. However more than the actual choice of the model parameters, we believe that the seminal contribution of this work is the creation of a comprehensive template for an upper limb model using which a large spectrum of models may be created corresponding to various healthy or pathological scenarios. The virtual physiology is neuromusculoskeletal in nature with a closed loop comprising motor efferents and proprioceptive afferents. A number of such virtual patients/physiologies may be created by choosing one or more model parameters from a probability distribution. This allows evaluating the effect of cell sizes, cell placements, orientations, synaptic weights, connection strategies, connection probabilities and many other chance factors on the movement of interest. Pathological physiologies may be created by reduction in cell numbers (atrophy) or controlled increase (neurogenesis), disruption of axons, tracts or loss of synaptic weights (injury, lesions), membrane properties (demyelination) modified connection rules (miswirings, genetic modifications), increased synaptic connections(increased excitability or sensitivity to muscle stretch), modification or removal of tendons, limbs (amputation or injury), modification of muscle or skeletal designs (prosthesis/implants), modified muscle lengths, insertion/origin points (surgical procedures) and the like. A number of motor programs may be simulated on the virtual patient by using appropriate spatio temporal patterns of stimulation of the various premotor interneurons and distal muscle motor neurons. This would be analogous to descending motor cortical activation of spinal cord motor programs [39]. Integration with premotor cortices, or BG models would be required in order to perform more complex and goal directed movements. This combined ability to create various movements in a large ensemble of healthy and pathological upper limbs we believe can pave the way for the arrival of virtual patient technology in movement disorders. The multiscale nature of the cervical spinal cord model makes it possible to create a variety of healthy and pathological conditions in this model including spinal cord injury, motor neuron diseases, spasticity, and lesions of the descending tracts to name a few. Together with musculoskeletal implants and prosthetics that can already be modeled in OpenSim, our work opens a virtual upper limb laboratory for in-silico tests and clinical trials of a broad spectrum of diseases. We hope that this work will open and facilitate in-silico medicine, experimentation, and clinical trials in movement disorders.

## V. Conclusion

We believe that this work constitutes the first report of a modular, multiscale neuro-musculoskeletal upper limb in-silico as a platform technology which can be used to create virtual patients. This technology can enable in-silico clinical trials by spawning large instances of physiologies that exhibit the variations commonly seen in populations in health and disease.

## Notes

### Competing Interest Statement

The authors have declared no competing interest.

