## Supplementary Text for "Neuro-musculoskeletal Upper Limb in-silico as virtual patient"

The supplementary text contains the information of every experiment, corresponding label of the behavioural outcome and the Gif of the movement generated. For the details of label (of Gifs) please check the catalogue section. The section contains labels for all the three models (Arm26, MOBL-Arm and Wrist).

Also note that for the open loop experiments involving MOBL-Arm the C4, C5 segment stimulation experiments have been aborted in between as the wrist joint reached the maximal range, in terms of radial deviation in case of C4 segment stimulation and wrist extension for C5 segment and the OpenSim integrator could not simulate further. So, the frames up to the halt position are alone uploaded in Gif’s folder.

Catalogue of Experiments: [Neuroid Experiments](https://docs.google.com/spreadsheets/d/e/2PACX-1vTZSGPC9NuOm0O1-Kd7EDHLt708vIjotdcyGd6F0IsTtheyoFkqJj3xG5Y8vOpuua2dgRkjHMiyo-xu/pubhtml)

To watch the behavioural responses in Gif: [Neuroid Gifs](https://drive.google.com/drive/folders/1kToA9-sNSAaI1m5H7nqkrRSmwq0weQeu?usp=sharing)
